# A 4-base pair core-enclosing helix in telomerase RNA is essential and binds to the TERT catalytic protein subunit

**DOI:** 10.1101/2020.01.10.902601

**Authors:** Melissa A. Mefford, Evan P. Hass, David C. Zappulla

## Abstract

The telomerase RNP counters the chromosome end-replication problem, completing genome replication to prevent cellular senescence in yeast, humans, and most other eukaryotes. The telomerase RNP core enzyme is composed of a dedicated RNA subunit and a reverse transcriptase (TERT). Although the majority of the 1157-nt *Saccharomyces cerevisiae* telomerase RNA, TLC1, is rapidly evolving, the central catalytic core is largely conserved, containing the template, template-boundary helix, pseudoknot, and core-enclosing helix (CEH). Here, we show that 4-base pairs of core-enclosing helix is required for telomerase to be active *in vitro* and to maintain yeast telomeres *in vivo*, whereas ΔCEH, 1-bp, and 2-bp alleles do not support telomerase function. Using the CRISPR/dCas9-based “CARRY two-hybrid” assay to assess binding of our CEH mutant RNAs to TERT, we find that the 4-bp CEH RNA binds to TERT, but the shorter-CEH constructs do not, consistent with the telomerase activity and *in vivo* complementation results. Thus, the CEH is essential in yeast telomerase RNA because it is needed to bind TERT to form the core RNP enzyme. Although the 8 nucleotides that form this 4-bp stem at the base of the CEH are nearly invariant among *Saccharomyces* species, our results with sequence-randomized and truncated-CEH helices strongly suggest that this binding interaction with TERT is dictated more by secondary than primary structure. In summary, we have mapped an essential binding site in telomerase RNA for TERT that is crucial to form the catalytic core of this biomedically important RNP enzyme.

## INTRODUCTION

Telomeres are repetitive sequences located at the ends of linear eukaryotic chromosomes. While they provide critical genome-protective functions, they are unable to be fully copied by DNA polymerases, owing to the end-replication problem. Short telomeres trigger a special G_2_/M cell-cycle arrest known as senescence. In order to overcome the end-replication problem and prevent senescence, most eukaryotic organisms require the ribonucleoprotein enzyme complex telomerase (Greider and Blackburn, 1985).

The telomerase core enzyme consists of a dedicated noncoding RNA subunit (TLC1 in *Saccharomyces cerevisiae*) and a reverse transcriptase protein component (TERT, or Est2 in *S. cerevisiae*). TERT utilizes a short template sequence in the telomerase RNA to iteratively add telomere repeats to the 3′ end of chromosomes (Greider and Blackburn, 1989). Together, these two core components are sufficient to reconstitute basal telomerase activity *in vitro* (Beattie *et al*., 1998; Zappulla *et al*., 2005). Telomerase RNAs are evolving strikingly fast, ranging in size from ~150 nucleotides in ciliates to >2000 nucleotides in some species of yeast, with even the sequences and secondary-structure models from closely related species within the same genus showing limited conservation. Experiments have shown that the 1157-nt *S. cerevisiae* telomerase RNA, TLC1, exhibits a high degree of organizational flexibility. First, TLC1 acts as a flexible scaffold to bind the holoenzyme proteins in the RNP enzyme complex: i.e., the binding sites for Est1, Ku, and Sm_7_ can each be repositioned to novel locations within the RNA while supporting these subunits’ functions (Zappulla and Cech, 2004; Zappulla *et al*., 2011; Mefford *et al*., 2013; Hass and Zappulla, 2017). Second, large portions of the RNA are dispensable for function *in vivo* and *in vitro* (Livengood *et al*., 2002; Zappulla *et al*., 2005; Qiao and Cech, 2008; Mefford *et al*., 2013).

Despite the wide array of differences between telomerase RNAs that have arisen during evolution, they do clearly share some key structural elements at their core (Lin et al. 2004). These include the (1) template, (2) template-boundary element, (3) pseudoknot with base triples, (4) core-enclosing helix (CEH), and (5) area of required connectivity (ARC).

In contrast to other conserved core elements, very little is known about the core-enclosing helix’s function. Being centrally located within the ARC, the CEH physically connects the pseudoknot to the template, enclosing the telomerase RNA’s core. However, when the CEH is deleted in the context of a functional circular permutation (i.e., maintaining RNA backbone integrity through the ARC), telomerase is inactivated *in vitro* (Mefford *et al*., 2013), suggesting a key role for the core-enclosing helix beyond simply enclosing the core. Consistent with the CEH being essential, large deletions that encompass either the 5′ or 3′ side of the core-enclosing helix cause senescence and disrupt Est2 binding *in vivo* (Livengood *et al*., 2002). Also suggesting that the CEH is a TERT-binding region, the CEH region has been shown to interact physically with a protein via gPAR-CLIP (Freeberg et al. 2013).

Here, we set out to investigate the structural and functional requirements of the core-enclosing helix in *S. cerevisiae*. We find that a core-enclosing helix of 4 base pairs is sufficient to provide telomerase function in yeast by promoting binding to TERT. There appear to be no sequence-specific requirements within the helix, indicating that TERT is generally binding double-stranded DNA. Together, these data convey the importance of the core-enclosing helix in yeast, while simultaneously explaining its sequence changes during evolution.

## RESULTS

### The core-enclosing helix is required to prevent senescence and support telomerase activity

Our previous results suggested that the core-enclosing helix is required for activity *in vitro* in the context of a circularly permuted telomerase RNA allele comprising just the catalytic core, Micro-T(170) (Mefford, Rafiq, and Zappulla 2013) (Fig. 1A). To investigate whether the core-enclosing helix is essential *in vivo* and to elucidate the structural requirements for function, we deleted or truncated the core-enclosing helix (CEH) in a circularly permuted larger telomerase RNA allele Mini-T(460). Switching of the context from Micro-T to Mini-T is necessary since Micro-T lacks features necessary for telomerase function *in vivo*. Using a circularly permuted RNA is necessary to avoid disrupting the Area of Required Connectivity when studying the CEH, which is in the center of the ARC (Mefford *et al*., 2013). Specifically, we chose the following two functional circular permutants: (1) cpJ3, which has the RNA ends between the template and the Est1 arm, and (2) cpTBE, which has the ends in the Ku-binding arm upstream of the template-boundary element (Mefford *et al*., 2013) (Fig. 1A and B).

**Figure 1.**
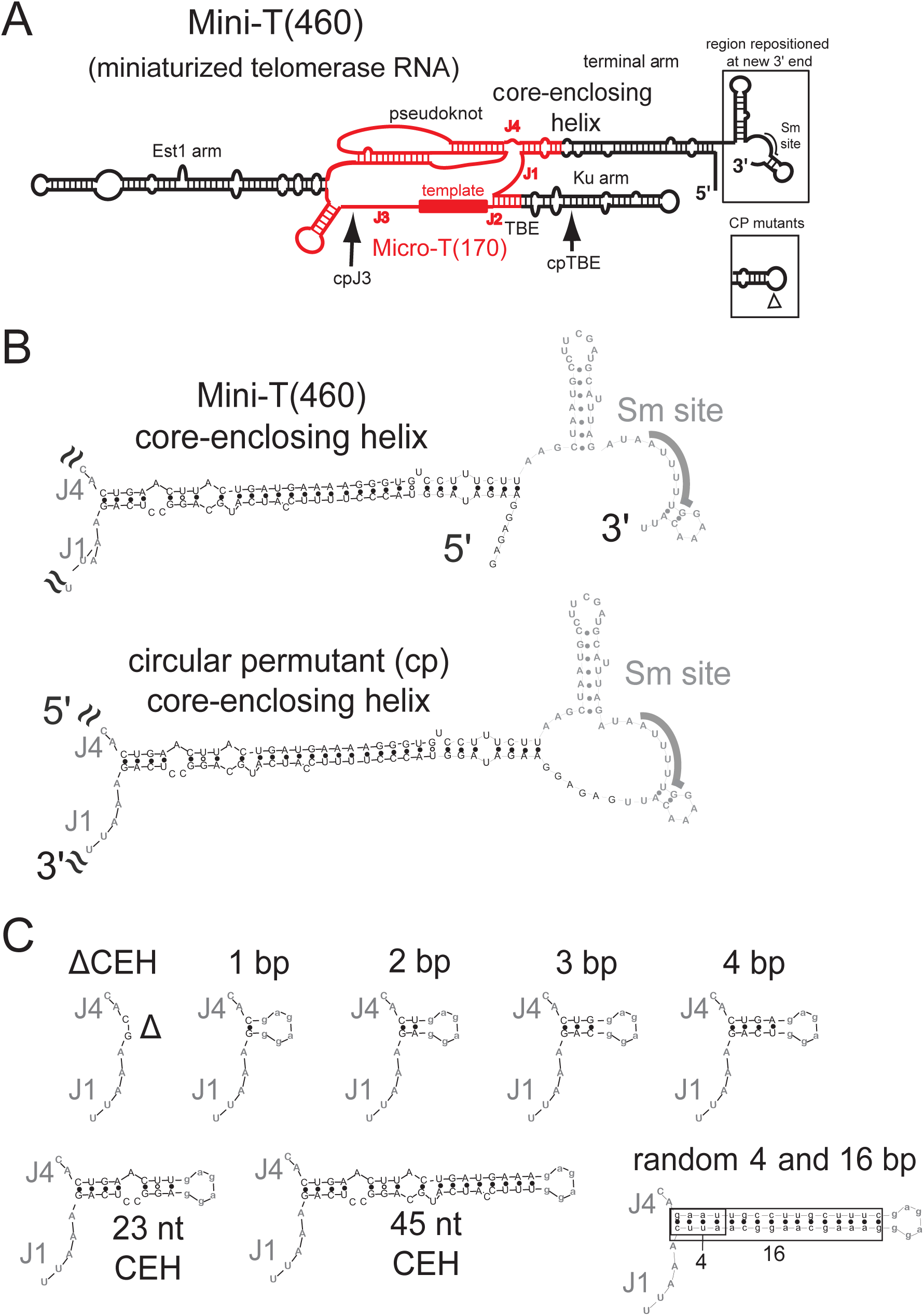
Core-enclosing helix mutations within circularly permuted Mini-T(460) yeast telomerase RNA. **A**. The Mini-T RNA is a smaller yet still functional version of the 1157-nt wild-type TLC1 RNA from *S. cerevisiae* (Zappulla et al., 2005). Shown in red is the Micro-T(170) RNA allele, which functions *in vitro* but not *in vivo*. The location of the repositioned 5′ and 3′ ends of the circular permutations cpJ3 and cpTBE are indicated by arrows. **B**. Circular permutation of Mini-T was performed by connecting the 5′ and 3′ ends of the processed RNA forms (accomplished by circularly permuting the gene encoding the RNA), shown below. **C**. Sequence of the core-enclosing helix truncations tested in Figs. 2 and 3.

We observed that the cpJ3 allele supported growth through 275 generations, as shown previously (Mefford *et al*., 2013), whereas the cpJ3ΔCEH allele (Fig. 1C) senesced by 75 generations, similarly to a Δ*tlc1* strain (Fig. 2A). Senescing cpJ3ΔCEH cells exhibited telomeres that were shorter than the cpJ3 cells on telomere Southern blots (Fig. 2B; lanes 13 and 14). The inability of cpJ3ΔCEH cells to maintain telomeres did not seem to be due to RNA abundance, since we detected nearly the same level of ΔCEH RNA by northern blotting as the functional cpJ3 allele (Fig. 2C). Furthermore, we observed that reconstituted telomerase (Zappulla *et al*., 2005) using cpJ3ΔCEH was catalytically dead *in vitro* (Fig. 2D). Thus, the core-enclosing helix is required for telomerase activity and function in cells.

**Figure 2.**
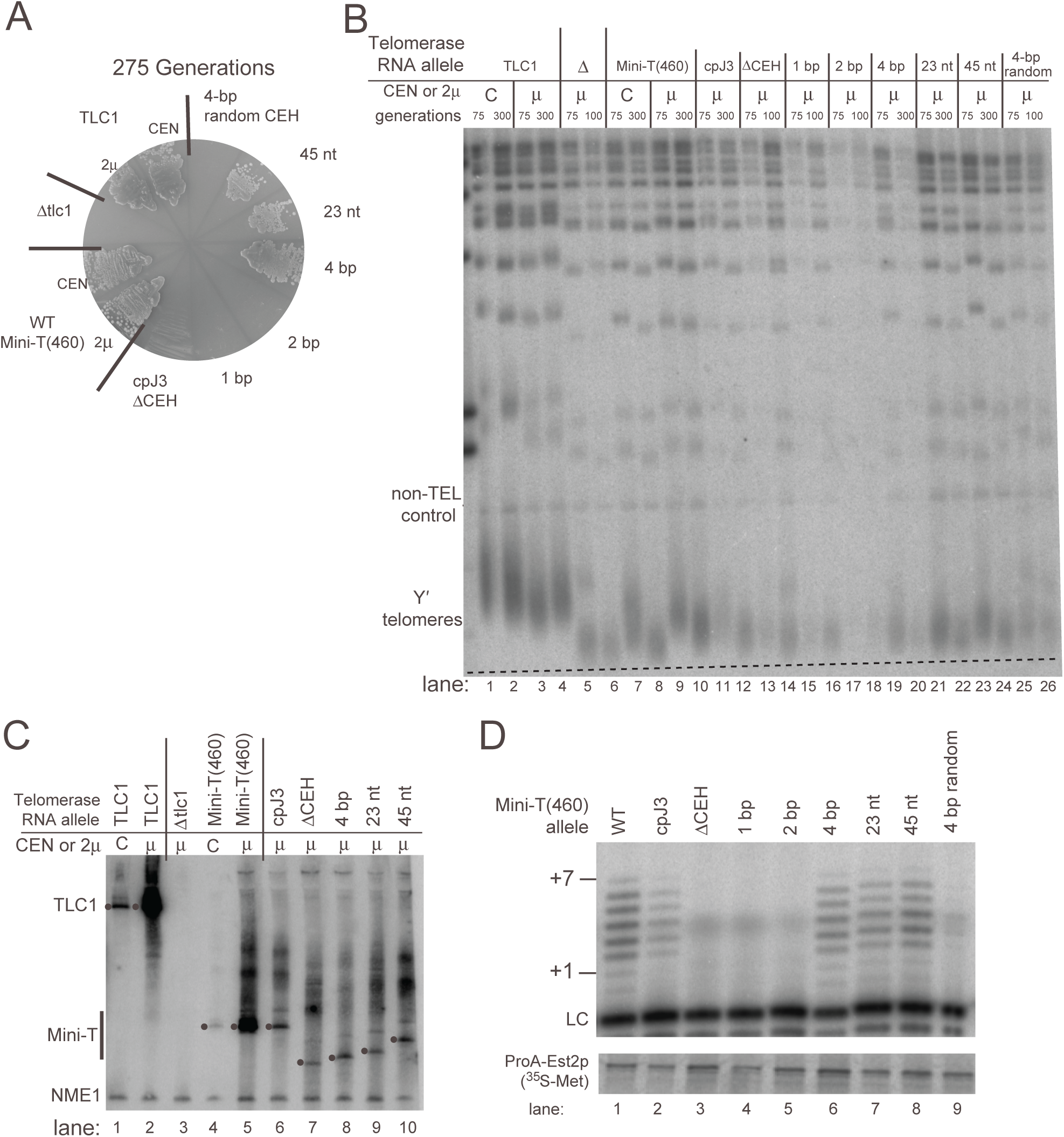
In Mini-T(460) cpJ3, a minimal core-enclosing helix of 4 base pairs is sufficient for telomerase activity *in vitro* and *in vivo*. **A**. *RAD52*-null yeast strains containing only the indicated telomerase RNA allele were serially restreaked through 275 generations. All cpJ3 core-enclosing helix variants were expressed from high-copy 2μ plasmids. The cpJ3ΔCEH, 1-bp, and 2-bp mutants senesce, while the 4-bp and longer mutants support growth. **B**. Viable cpJ3 core-enclosing helix mutants maintain short telomeres. Separated *XhoI*-digested genomic DNA was probed by Southern blotting for telomeric repeat sequence and a chromosome IV control region. Note that the number of generations yeast were grown vary as indicated depending on their viability in liquid culture. The dotted line represents the average length of Y′ telomeres in Mini-T(460) at 300 generations when expressed from a 2μ plasmid. The slant of the line reflects the migration of the Ch. IV control band. **C**. Mature telomerase RNA is detectable from core-enclosing helix variants. Total cellular RNA was probed for either a fragment corresponding to Mini-T(460) or a 340-nt control RNA, NME1. The black dots to the left of bands indicate the full-length RNA, which differ in length according to the amount of core-enclosing helix sequence present. **D**. Only viable core-enclosing helix mutants show detectable *in vitro* telomerase activity. Radiolabeled products of telomerase reactions (+1 to +7) were separated on a 10% acrylamide denaturing gel by electrophoresis. A recovery and loading control (LC) was included in the telomerase reactions. Below, TERT protein containing ^35^S-methionine was separated by denaturing protein gel electrophoresis to show immunopurification of telomerase from the *in vitro* transcription and translation system.

To further test the conclusion that the core-enclosing helix is essential in yeast telomerase, we examined telomerase function when the CEH was deleted in a different circular permutant, cpTBE. As in the context of cpJ3, we observed that cpTBEΔCEH senesced by 75 generations (Fig. 3A) concomitant with shortening telomeres observed by Southern blotting (Fig. 3B, lane 12). Senescence was again not due to lack of RNA accumulation, as the cpTBEΔCEH transcript was evident by northern blotting at levels similar to other functional alleles (Fig. 3C, lane 7). Furthermore, the Mini-T(460) cpTBEΔCEH allele was catalytically dead *in vitro* (Fig. 3D, lane 3).

**Figure 3.**
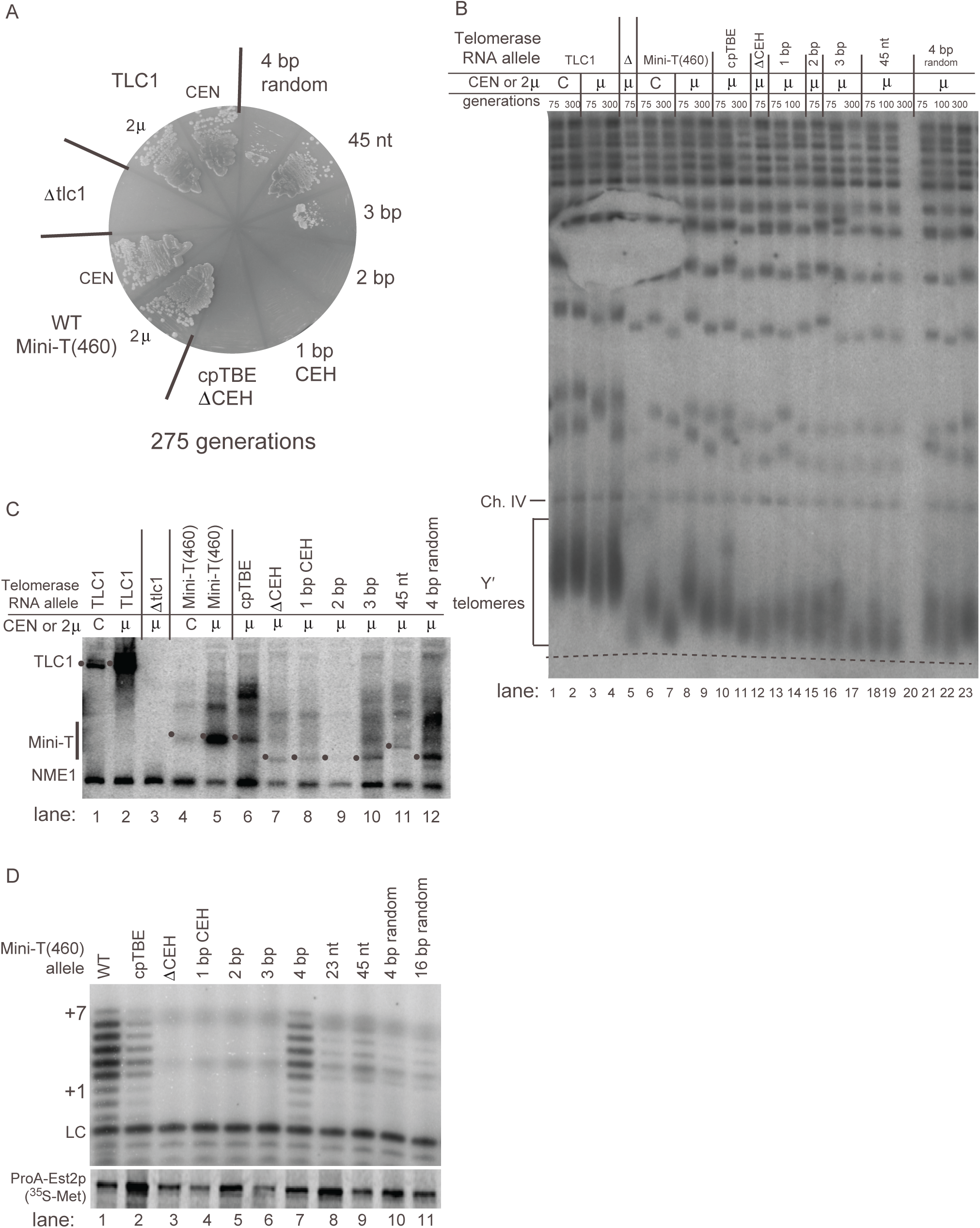
In Mini-T(460) cpTBE, a minimal core-enclosing helix of 3 base-pairs prevents senesce and 4 base-pairs supports robust enzyme activity *in vitro*. **A**. Yeast expressing the Mini-T(460) 3 bp allele show stable small colony growth on plates. Yeast were serially restreaked as in Figure 1. All core-enclosing helix variants were expressed from 2μ plasmids. **B**. Viable core-enclosing helix mutants maintain telomeres, though they are shorter than wild-type Mini-T(460). Southern blot was performed as in Figure 1, with a dotted line passing through wild-type 2μ Mini-T(460) at 300 generations. Some yeast could not be grown in liquid culture to the 300-generation time point as indicated. **C**. Mature telomerase RNA is detectable for all core-enclosing helix mutants. Northern blotting was performed as in Figure 1. The location of the mature telomerase RNA is indicated by a black dot to the left of each band, depending on expected size. **D**. Telomerase activity *in vitro* is barely detectable for cpTBE 3 bp, while mutants with 4 base pairs or more of core-enclosing helix show detectable telomerase activity. *In vitro* telomerase activity assays were performed as in Figure 1, with immunopurified Est2p below.

Together, the independent results from the contexts of both the cpJ3 and cpTBE circular permutations show that the core-enclosing helix is essential for telomerase function *in vivo* and RNP core-enzyme activity *in vitro*.

### A core-enclosing helix of 4 base pairs is sufficient to support telomerase activity

Given the functional necessity of the core-enclosing helix, we set out to determine what features of its structure are required. We began by adding back the native core-proximal base pairs one by one (Fig. 1C). This analysis revealed that just 4-base pairs of native core-enclosing helix sequence in cpJ3 supported telomerase function in preventing senescence (Fig. 2B) and telomerase activity with TERT *in vitro* (Fig. 2D). A CEH of 3 base pairs in cpTBE caused an intermediate phenotype, barely allowing cells to avoid senescence (and causing slow growth of those that did survive) (Fig. 3A), and without clearly perceptible telomerase activity (Fig. 3D, lane 6). In contrast, 1- and 2-bp core-enclosing helices in cpJ3 did not support any telomerase function *in vitro* or *in vivo* (Fig. 2). For these alleles, it is likely that base pairing does not stably form. Thus, our data show that a minimum of 4 base pairs is required for substantial core-enclosing helix function.

Adding back more of the native core-enclosing helix also supported telomerase activity *in vivo* and *in vitro*, though with no detectable increases in activity over the shorter 4-base pair helix (Figs. 2 and 3). Specifically, we added back either 23 nucleotides, which corresponds to the endogenous core-enclosing helix present in Micro-T(170), or 45 nucleotides. Both of these core-enclosing helices supported cell growth without evidence of senescence in the two circularly permuted Mini-T contexts (Figs. 2A and 3A), stably short telomeres (Fig. 2B, lanes 21–24; Fig. 3B, lanes18–20), and reconstituted telomerase activity (Fig. 2D, lanes 7 and 8; Fig. 3D, lanes 8 and 9).

### The native core-proximal sequence of the core-enclosing helix is not necessary for function

To determine whether the sequence of the native core-enclosing helix sequence is important, we tested non-native base pairs. Interestingly, a sequence-randomized 4-bp helix (Fig. 1C; GAAU vs. wild-type CUGA) was able to support weak telomerase activity in the context of cpTBE, both *in vivo* (Fig. 3A; Fig. 3B, lanes 21–23) and *in vitro* (Fig. 3D, lane 10). The same random-sequence 4-bp helix in the context of cpJ3 was not sufficiently functional *in vivo* to prevent senescence (Fig. 2A), although it had weakly perceptible activity *in vitro* (Fig. 2D). A longer 16-bp random helix (Fig. 1C) in cpTBE showed similar telomerase function to the 4-bp random helix (Fig. 3D). These results suggested that the endogenous sequence of the core-enclosing helix has a nonessential role in its function.

To further test the structural and functional requirements of the core-enclosing helix, we examined a series of deletions in non-circularly permuted Mini-T(460). If the particular sequence or the positioning of bulges in the core-enclosing helix are crucial for function, then deleting these regions should abolish function. We first deleted the four base pairs nearest to the core (Δ8; Fig. 4A). These are the same four base pairs that supported function in cpJ3 (Fig. 2). This Δ8 CEH-truncation allele causes the core-enclosing helix to now begin at an asymmetrical 3-nt bulge, increasing the length of junctions J4 and J1 on either side of the helix, followed by 3 base pairs (Fig. 4A). This Δ8 deletion supported growth through 275 generations (Fig. 4B), although telomeres in these mutants were shorter than those of wild-type Mini-T(460) cells (Fig. 4C, lanes 8 and 9 versus lanes 10 and 11). The shorter telomeres in Mini-T(460)Δ8 do not appear to be due to lower RNA (Fig. 4D, lane 5 versus lane 6). However, the deletion of 8 nucleotides did partially decrease telomerase core-enzyme activity *in vitro* (Fig. 4E, lane 1 versus lane 2).

**Figure 4.**
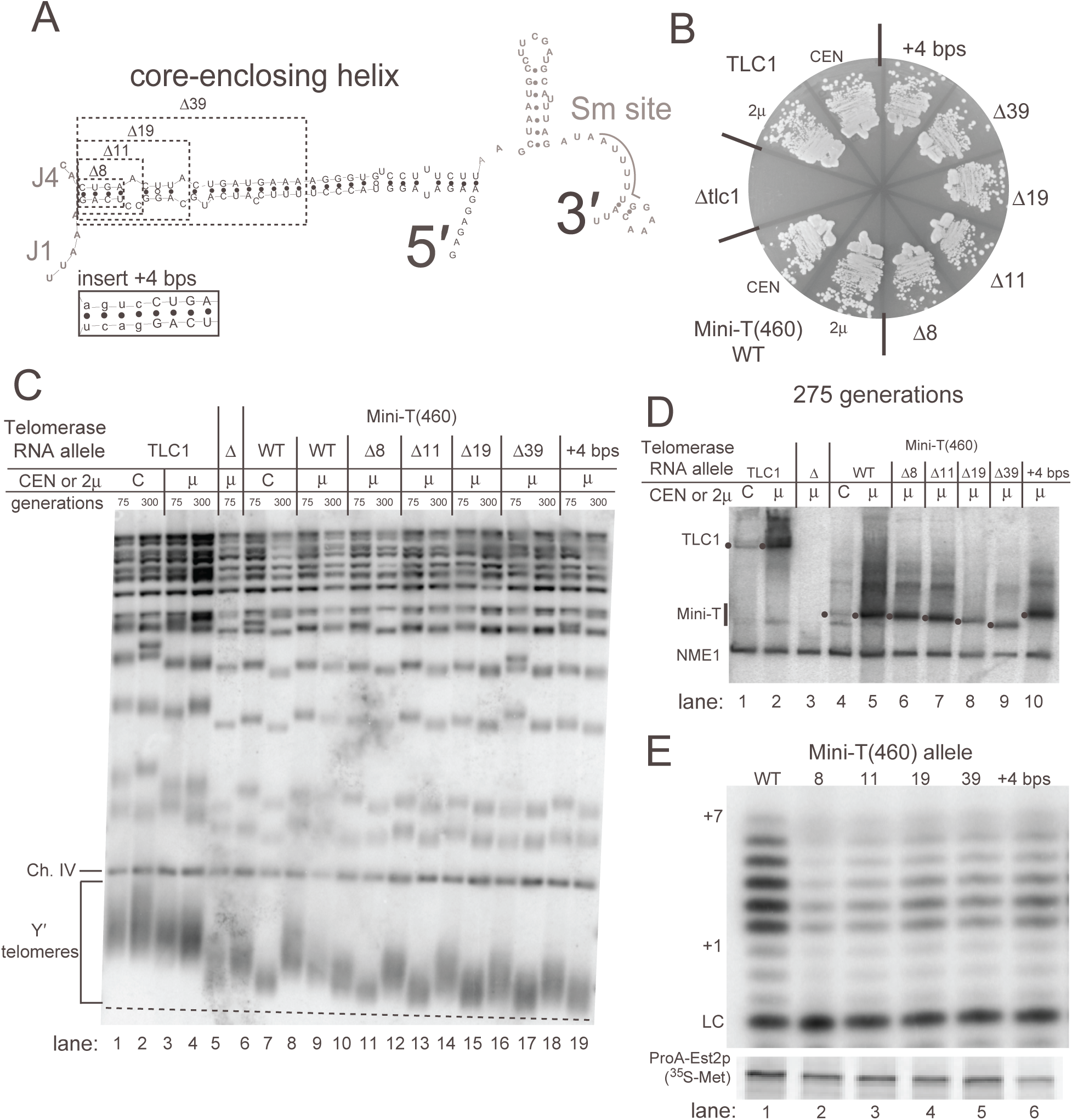
The core-proximal regions of the core-enclosing helix are dispensable for telomerase function *in vitro* and *in vivo*. **A**. Schematic indicating the native core-enclosing helix nucleotide deletions tested. Inset shows the sequence of four base pairs inserted on the core-proximal side in the +4 bp allele. **B**. Mutations that alter the length and nature of the core-proximal region of the core-enclosing helix prevent senescence in Mini-T(460). Yeast were restreaked on solid media through 275 generations. All deletion or insertion mutants were expressed from high-copy 2μ plasmids. **C**. Telomeres are shorter relative to Mini-T(460) when the core-enclosing helix is truncated or extended. Genomic DNA was Southern blotted as in Figure 1. The dotted line represents the length of wild-type Mini-T(460) telomeres at 300 generations for comparison. **D**. Telomerase RNA is expressed from all mutants tested. Northern blots were performed as in Figure 1 with black dots indicated the position of the mature RNA in the gel. **E**. Deletions and insertion cause slight defects in telomerase activity *in vitro*. Telomerase activity assays were completed as in Figure 1. Immunopurified radiolabeled Pro-A-Est2 protein is shown below.

Larger deletions of the core-enclosing helix also supported telomerase activity. We deleted 11 nucleotides (corresponding to the native nucleotides present in Micro-T(170)), 19 nucleotides, and 39 nucleotides (Fig. 4A). All of these deletions result in different core-enclosing helical structures near the core, yet each of these alleles supported telomerase activity *in vivo* and *in vitro* (Fig. 4B, E). The shortened telomeres (Fig. 4C) supported by these alleles do not appear to be due to insufficient levels of the RNAs (Fig. 4D).

Finally, we tested insertion of an additional 4 base pairs on the core-proximal end of the core-enclosing helix (Fig. 4A). This insertion (+4 bp) prevented senescence through 275 generations (Fig. 4B). Similar to the truncations of the CEH, the +4-bp insertion also caused telomere shortening relative to wild-type Mini-T(460) (Fig. 4C, lanes 18 and 19 versus lanes 8 and 9), not attributable to lowered telomerase RNA expression (Fig. 4D, lane 5 versus lane 10). The +4-bp insertion also supported telomerase activity *in vitro* (Fig. 4E).

### The core-enclosing helix is required for telomerase RNA binding to TERT

What is the mechanistic role of the core-enclosing helix in telomerase RNP function? Our results show that the CEH is required for both *in vivo* and *in vitro* telomerase action — thus, the CEH is essential for fundamental telomerase core-enzyme activity. Since the core enzyme is composed of the TLC1 RNA and the TERT catalytic protein subunit, a parsimonious explanation for the contribution of CEH to enzyme function is that it is simply required for the RNA to bind to TERT to assemble the core enzyme.

To test the hypothesis that the CEH is a key binding site for TERT in telomerase RNA, we used the *in vivo* CRISPR-assisted RNA-RBP yeast (CARRY) two-hybrid RNA-protein interaction assay (Hass and Zappulla, *bioRxiv* 2017). This method employs catalytically inactive CRISPR/dCas9 to tether an RNA of interest upstream of reporter genes in yeast such that if the RNA of interest binds to a protein (fused to a Gal4 transcriptional-activation domain; GAD), the RNA-protein binding interaction can be detected via *HIS3* reporter-gene expression, allowing yeast to grow in the absence of histidine (Figure 5A) (Hass and Zappulla, *bioRxiv* 2017). First, we tested if Micro-T(170) binds to TERT in the CARRY two-hybrid system. Indeed, the Micro-T(170) construct allowed significant growth on media lacking histidine (~1000-fold greater than the negative control; Fig. 5B), indicating strong reporter-gene activation based on binding to GAD-Est2 (TERT). Furthermore, circularly permuted Micro-T (cpTBE) also bound at least as well to GAD-TERT as unpermuted wild-type Micro-T. However, when we deleted the CEH from this cpTBE allele, there was a complete loss of TERT binding (Fig. 5B, compare rows 3 and 4). Micro-T cpTBE with a 1-bp, 2-bp, or 3-bp CEH did not bind TERT, but a 4-bp CEH did, consistent with performance of these alleles in telomerase function *in vivo* and *in vitro* (Figs. 2 and 3). We also tested binding by the random 4-bp and 23-nt CEH alleles and these showed little or no HIS activation. Presumably these alleles bind intermediately to TERT; too weakly for the CARRY two-hybrid system to detect (≥ ~1 µM K_d_; (Hass and Zappulla, 2017)), yet still sufficient to provide telomerase assembly and function, as shown in Figure 2 and 3. Overall, we conclude that the CEH is required to bind to TERT and that 4 base pairs is sufficient for this critical interaction at the core of the telomerase RNP.

**Figure 5.**
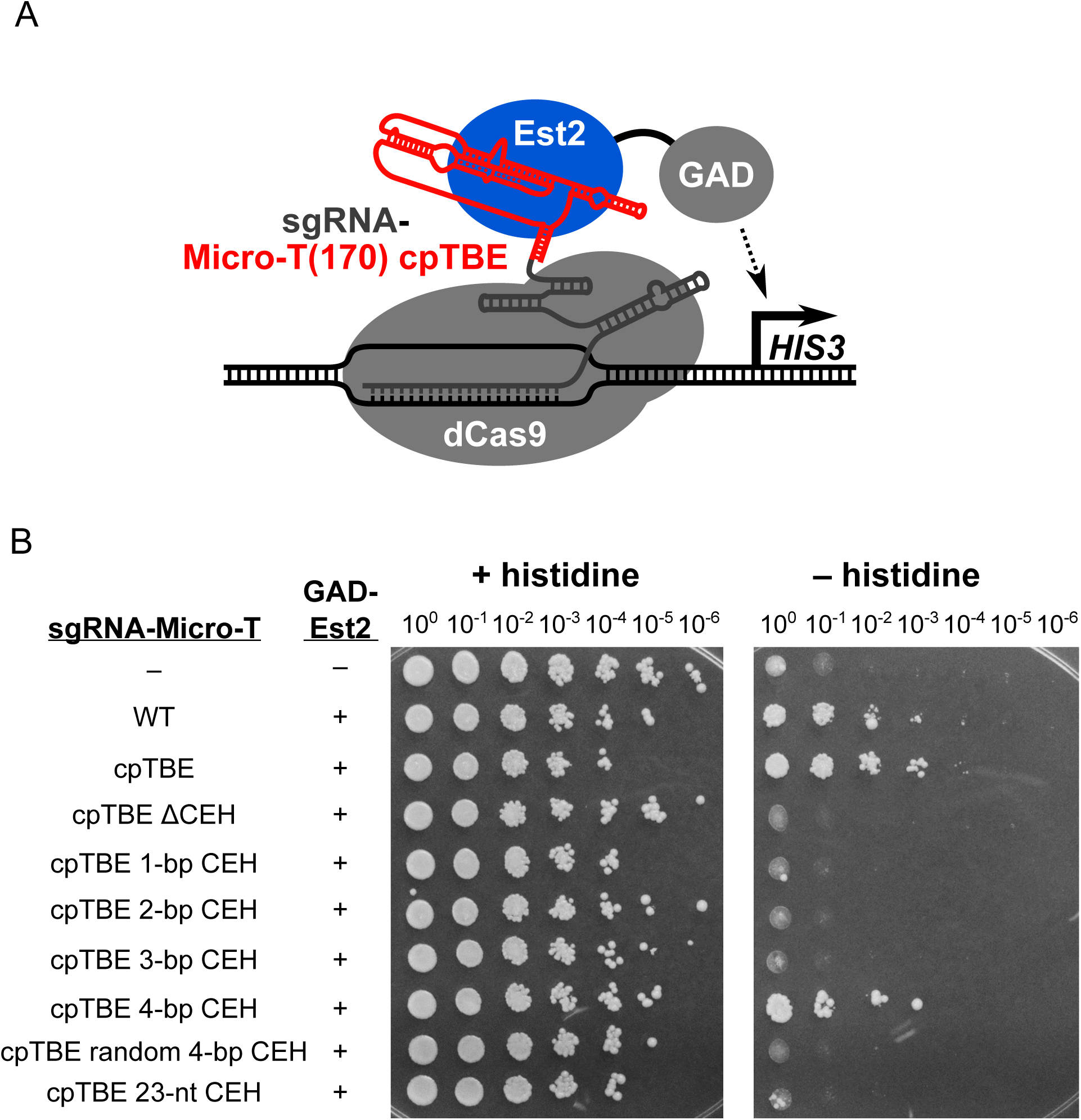
TERT-binding to telomerase RNA is impaired by a core-enclosing helix shorter than 4 base pairs. **A**. Schematic of the CRISPR-assisted RNA-RBP yeast (CARRY) two-hybrid system (Hass and Zappulla, 2017). Different mutants of the TLC1 RNA core were fused to a CRISPR sgRNA which was tethered to the promoter of a *HIS3* reporter gene by nuclease-dead Cas9 (dCas9). A fusion of Est2 to the Gal4 transcriptional activation domain (GAD) was also expressed, and activation of the *HIS3* reporter gene was used as a measure of binding between Est2 and the TLC1 RNA core. **B**. A 4-bp core-enclosing helix is required for Micro-T(170) binding to TERT in the CARRY two-hybrid system. Expression of *HIS3* was assayed by growing cells to saturation in liquid culture, making six 10-fold serial dilutions, and spotting these cells to solid media with or without histidine. In the GAD-Est2 column on the left, “-” indicates expression of GAD that is not fused to Est2 while “+” indicates expression of the GAD-Est2 fusion protein. Similarly, “–” in the sgRNA-Micro-T column indicates expression of an sgRNA that is not fused to Micro-T.

## DISCUSSION

A multi-subunit enzyme cannot be active if its core components do not assemble. Despite this basic criterion for function, little is known about the interfaces that govern binding between the subunits at the heart of the biomedically relevant telomerase RNP enzyme. Here we have demonstrated that the core-enclosing helix of telomerase RNA is essential for binding to the TERT catalytic protein subunit in yeast.

In order to study the core-enclosing helix, which is in the middle of the Area of Required Connectivity, we moved the ends of telomerase RNA away from the ARC so as not to disrupt it in our CEH-mutant alleles. Thus, we circularly permuted TLC1 RNA, allowing us to delete the CEH without disrupting RNA continuity through the ARC. The ability to parse ARC from CEH function revealed that the CEH is essential for telomerase RNP function both *in vivo* and *in vitro*, consistent with it being required for binding to TERT. Furthermore, our findings are based on data obtained from studying the CEH within two independent circularly permuted telomerase RNA allele contexts (Figures 2 and 3). One of the circular permutations has the RNA ends moved to the distal portion of the template-boundary element, cpTBE, and the other to within the junction in the central hub between the template and pseudoknot, cpJ3 (Figure 1A). Under both of these different conditions, we found that a 4-bp CEH is sufficient for its function. Fewer base pairs did not permit CEH function, and it is likely that they also do not stably form a helix, providing additional evidence that a paired secondary structure element is necessary at this position within the catalytic core of yeast telomerase RNA.

As for any functional importance of the sequence of the CEH, our results from sequence-randomized, truncated, and extended-helix alleles provide strong evidence that core-enclosing helices with diverse sequences (6 different variations) provide at least basic telomerase activity. Functionality of the CEH in these alleles was despite the sensitized context of being tested within miniaturized TLC1 RNA, which, even when otherwise wild-type in sequence, supports rather short telomeres.

The core-enclosing helix is a conserved feature of telomerase RNAs beyond yeasts, being also present in human, ciliate, and most other telomerase RNAs known to date. Although the CEH 4-base pairs that we find to be critical in *S. cerevisiae* are invariant among *Saccharomyces* (Dandjinou *et al*., 2004; Zappulla and Cech, 2004), evidence suggests that core-enclosing helices are not absolutely required in all known telomerase RNAs. For example, disrupting the core-enclosing helix in *Tetrahymena thermophila* reduced activity by 95–99%, thus not quite abolishing it altogether (Mason *et al*., 2003). In human telomerase RNA, deleting the 5′-end still permits functional telomerase, although template boundary definition is compromised (Chen and Greider, 2003). Curiously, rodent telomerase RNAs have the 5′ end located just 2–5 nucleotides upstream of the template, and presumably do not form a core-enclosing RNA helix (Hinkley *et al*., 1998; Chen *et al*., 2000). In these species, it has been proposed that TERT simply uses run-off reverse transcription, since these RNAs also lack a template-boundary element (Hinkley *et al*., 1998), so thus differ from yeast and humans. It is worth also noting that the template-boundary element and core-enclosing helix appear to be consolidated into a single paired element in human telomerase RNA (Chen and Greider, 2003; Lin *et al*., 2004). Thus, the CEH in yeast could be an essential TERT-binding site while not performing the orthologous function in some other species. It is likely that the reason for differences with respect to how the RNA and TERT interact is due to the rapid evolution of the RNP.

Finally, our results also show that binding of TERT to TLC1 is not helix sequence-specific, but rather is dictated by secondary structure. Six different paired sequences were tested in place of the CEH, all of which functioned. Overall, our data suggest that TERT binds directly to the core-enclosing helix. Nevertheless, it is feasible that the CEH promotes binding of TERT elsewhere in the RNA. As for which of the other conserved secondary-structure elements in yeast telomerase RNA might also be binding TERT, some possibilities include the pseudoknot, template-boundary element (TBE), and core junctions. The *Tetrahymena* TERT RNA-binding domain makes contacts with nucleotides at the base of the TBE helix (Jansson *et al*., 2015), and it is possible that the yeast telomerase TBE and/or CEH make similar binding interactions with Est2.

Telomerase is a unique reverse transcriptase in that it, unlike viral reverse transcriptases, iteratively re-uses a short RNA template intrinsic to the RNP enzyme. This template re-use required for processive activity involves transient disruption of the base pairing of the RNA with the enzyme’s DNA substrate while the RNP remains associated with the substrate via another substrate-enzyme binding interaction. During this multifaceted interaction of the enzyme with telomeric DNA, the telomerase RNA-TERT interaction must also be maintained. This highly dynamic interplay should necessitate a multi-partite binding interaction between the telomerase RNA and TERT. Telomerase of course interacts with the template via its active site, but we hypothesize that the highest-affinity binding interaction between the RNA and TERT resides outside of the RNA’s template region in order to provide a foothold for maintaining core RNP integrity during catalysis. Such a template-distal TERT-binding site in the RNA may be important to avoid sterically hindering the repeated process of binding, extending, and dissociating from the telomeric DNA substrate. Specifically, we propose that TERT binds most prominently to telomerase RNA nucleotides elsewhere around the RNA’s central hub from the template in the secondary structure model (Fig. 6). This is supported by our current results as well as our prior finding that breaking the RNA backbone in the Area of Required Connectivity (ARC), which connects the core-enclosing helix to the template, abolishes telomerase core enzyme activity (Mefford *et al*., 2013). Furthermore, prior molecular-genetic studies in yeast have provided evidence that key binding site(s) for TERT in TLC1 reside in this vicinity (Livengood *et al*., 2002; Chappell and Lundblad, 2004; Qiao and Cech, 2008; Freeberg *et al*., 2013; Mefford *et al*., 2013).

**Figure 6.**
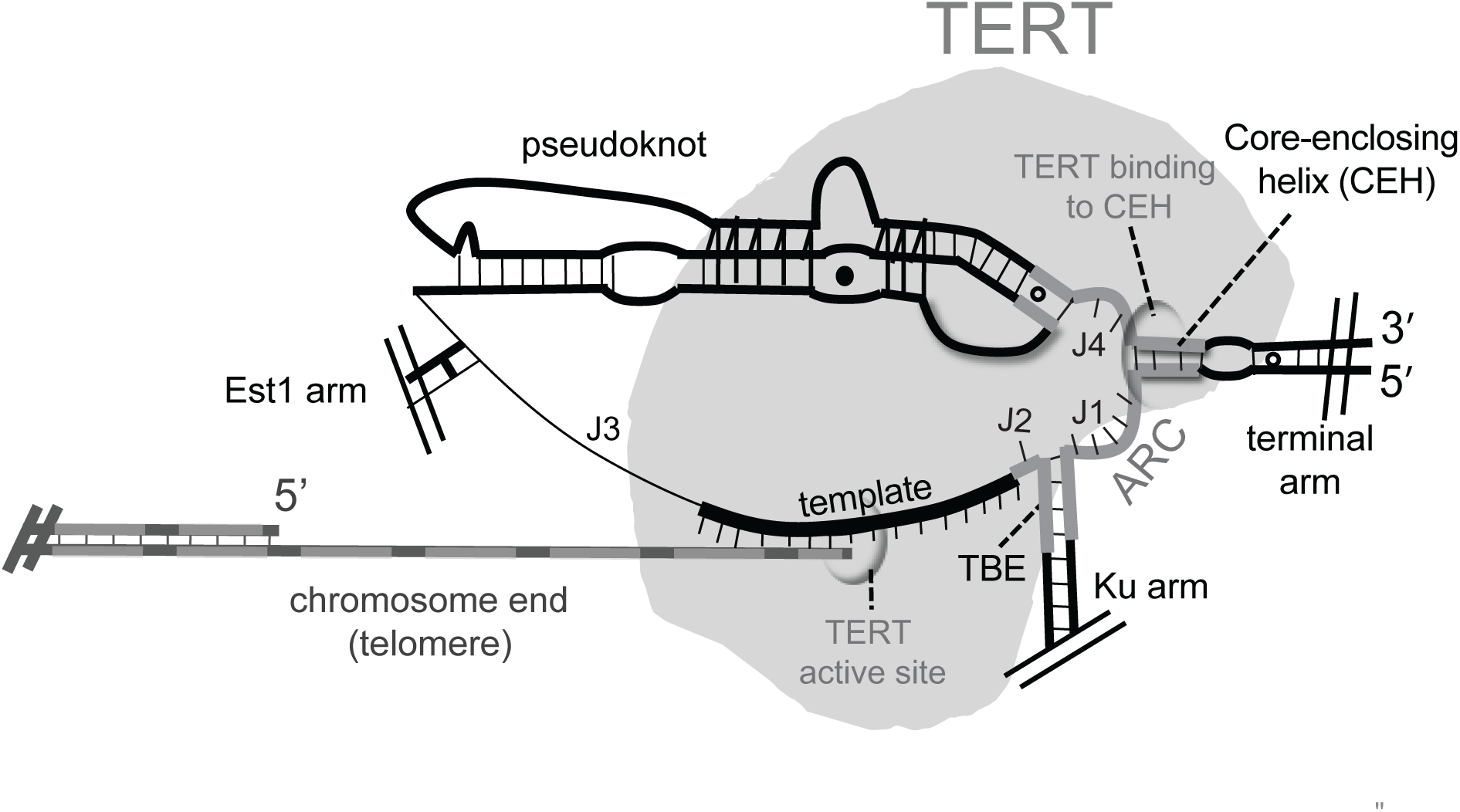
RNA-TERT interactions and their functional implications for telomere extension by the yeast telomerase RNP core enzyme. The core-enclosing helix and its binding to TERT (gray) is indicated. We hypothesize that binding of TERT to region(s) connected to the template by the Area of Required Connectivity (dark gray) allow TERT to stay associated with TLC1. This binding interaction is likely to be important during the unique telomerase reverse-transcription catalytic cycle that is based on an intrinsic RNA template, which requires the TERT active site to dissociate from the template and newly synthesized DNA during repeat-addition processivity *in vivo*.

## MATERIALS AND METHODS

### Yeast strains and plasmids

All *in vivo* assays for telomerase function used yeast strain yVL009 (*MATa tlc1Δ::LEU2 rad52-Δ::LYS ura3-52 lys2-801 ade2-101 trp-1-Δ1 his3-Δ200 leu2-Δ cir*^*+*^ pSD120/p*TLC1*-*URA3*-*CEN*) (Chappell and Lundblad, 2004). All new mutants were made by PCR and subcloned into a 2μ *TRP1*-marked pRS424 vector for *in vivo* assays, or a pUC19-based vector for *in vitro* assays.

### Senescence assays

yVL1009 yeast were transformed with a *TRP1*-marked *CEN* or 2μ plasmid via standard lithium acetate methods. Transformants were selected on media lacking uracil and tryptophan, and then streaked to media lacking tryptophan and containing 5-FOA to counter-select for the wild-type *URA3*-marked TLC1 cover plasmid. Duplicate colonies of yeast containing only the transformed plasmid were then serially streaked ten times on media lacking tryptophan at 30°C.

### Southern blots

To determine telomere length, Southern blots were performed as previously described (Zappulla *et al*., 2011). Briefly, yeast cells were cultured in liquid media lacking tryptophan at 30°C, and genomic DNA was isolated (Puregene kit, Qiagen). Purified genomic DNA was digested overnight with XhoI, then electrophoresed through a 1.1% agarose gel at 70V for 17 hours. DNA fragments were then transferred to a Hybond N^+^ membrane (GE Healthcare) by capillary action, crosslinked with UV irradiation, and incubated with radiolabeled probes for telomeric DNA and a region of chromosome IV. Blots were imaged using a Typhoon 9410 Variable Mode Imager.

### Northern blots

Total RNA was isolated, using a hot-phenol extraction method, from log-phase yeast liquid cultures grown in the absence of tryptophan. 30 μg of total cellular RNA was electrophoresed through a 4% polyacrylamide/1X TBE/7M urea gel at 35 W for ~3 hours. Separated RNAs were electro-transferred to a Hybond N^+^ membrane (GE Healthcare) and UV crosslinked. The membrane was probed with 1 × 10^7^ cpm of a StuI-NsiI fragment of Mini-T(460) and 1 × 10^5^ cpm of the control RNA, *NMEI*. Blots were imaged using a Typhoon 9410 Variable Mode Imager.

### Reconstituted telomerase activity assays

Telomerase was made using a coupled *in vitro* transcription and translation system as previously described (Zappulla *et al*., 2005). Briefly, DNA templates for ProA-Est2 TERT and telomerase RNA were incubated in a rabbit reticulocyte lysate system including ^35^S-methionine for 90 minutes. Telomerase was then immunopurified using IgG Sepharose beads (GE Healthcare). Purified telomerase was incubated with telomeric DNA oligo (DZ428, 5′-GGTGTGGTGTGGG-3′), a 5′ *γ*^32^P labeled internal recovery and loading control, *α*^32^P dTTP, and 1 mM each unlabeled dATP, dCTP, dGTP in standard *in vitro* telomerase activity buffer. Assembled reactions were incubated for 10 minutes at 26°C, then stopped by the addition of ammonium acetate and ethanol precipitated. Products were resuspended in formamide loading dye and boiled at 95°C for 5 minutes. Samples were electrophoresed through a 10% polyacrylamide/1X TBE/7M urea gel for 75 minutes at 90 W and then dried on Whatman paper and exposed to a phosphorimager screen. Imaging was done with a Typhoon 9410 Variable Mode Imager. As a loading and recovery control during purification of reconstituted telomerase (Figs. 2D, 3D, and 4E), 2.5 ml of telomerase was resuspended in Laemmli sample buffer, boiled at 95°C for 5 minutes, and separated on a 7.5% Mini-PROTEAN TGX gel (BioRad). The gel was transferred to Whatman paper, dried, and imaged using a Typhoon 9410 Variable Mode Imager.

### CARRY two-hybrid RNA-protein binding assays

CARRY two-hybrid assays were carried out as described previously (Hass and Zappulla, 2017). Briefly, the strain CARRYeast-1a (*MATa his3Δ200 trp1-901 ade2 lys2::(4LexAop-HIS3) ura3::(8LexAop-LacZ) leu2::(KanMX6_dCas9)*) was transformed with sgRNA-fusion and GAD-fusion plasmids with the vector backbones from *TRP1*-marked pCARRY2 (Hass and Zappulla, 2017) and *LEU2*-marked pGAD424 (Bartel *et al*., 1993), respectively. Transformants were grown to saturation in synthetic liquid culture medium lacking tryptophan and leucine. Six 10-fold serial dilutions were made of each culture, and 5 μL of each dilution, as well as the undiluted culture, were pipetted onto both solid –Trp –Leu and –Trp –Leu –His minimal media. Cells were incubated for 3 days at 30°C and photographed.

## ACKNOWLEDGEMENTS

Research reported in this publication was supported by the National Institute of General Medical Sciences of the National Institutes of Health under award number R01GM118757 to D.C.Z.

